# Genetic control of neuronal activity enhances axonal growth only on permissive substrates

**DOI:** 10.1101/2022.01.21.477184

**Authors:** Francina Mesquida-Veny, Sara Martinez-Torres, José Antonio Del Río, Arnau Hervera

## Abstract

Neural tissue has limited regenerative ability, to cope with that, in the recent years a diverse set of novel tools have been used to tailor neurostimulation therapies and promote functional regeneration after axonal injuries. In this report, we explore cell-specific methods to modulate neuronal activity, including opto- and chemogenetics to assess the effect of specific neuronal stimulation in the promotion of axonal regeneration after injury. We found that opto- or chemogenetic modulations of neuronal activity on both dorsal root ganglia and corticospinal motor neurons increase their axonal growth capacity only on permissive substrates.

## Background

Following injury, damaged axons from the central nervous system (CNS) degenerate and are unable to regenerate, while surviving fibres, have a limited capacity to sprout and to re-establish lost connections, leading to functional impairment [1].

This failure of CNS axons to regenerate after injury is partly attributed to a hostile CNS environment for growth [2–4]. The injury site is rich in growth inhibitory proteins including myelin-associated glycoproteins, and chondroitin sulfate proteoglycans (CSPG) while lacking trophic support for axon regeneration [5–7]. The limited intrinsic regenerative capacity of adult CNS axons also contributes to the failure of axon regeneration after injury [8–10]. Yet, in the last years there have been several studies approaching different aspects of the CNS physiology to both boost the intrinsic regrowth capacity and overcome the extrinsic inhibition [11, 12].

During development, when CNS neurons still retain their intrinsic ability to regenerate after axotomy [13], neuronal activity critically determines their connectivity, particularly onto spinal targets [14]. In this direction, neuronal activation has also been explored to try to increase the intrinsic capacity for axon regeneration as well as overcome the extrinsic inhibition. Practically, electrical stimulation has been shown to enhance regeneration of sensory axons after peripheral nerve or dorsal columns injury [15, 16] and sprouting of cortical axons into contralateral spinal cord gray matter after pyramidotomy [17–19].

In order to gain insight on the actual mechanisms underlying these improvements, specifically and remotely activating neurons through opto- and chemogenetics, has become the state-of-the-art approach for tailored activity modulation [20, 21].

In this direction, the current study explores the therapeutic potential of modulating neuronal activity of sensory (dorsal root ganglia (DRG)) and motor descending (corticospinal motor neurons (CSMN)) neurons, via opto- and chemogenetic stimulation in regulating axonal growth after injury.

We found that specific neuronal stimulation induced increased axonal growth capacity both *in vitro* and *in vivo*, and both in sensory and motor neurons, but only when in presence of permissive and trophic environments. Opto- or chemogenetic stimulation did not overcome the inhibitory signalling induced by molecules such as myelin-associated glycoproteins and CSPGs.

## Methods

### Mice

B6.Cg-Tg(Thy1-COP4/EYFP)18Gfng/J (Thy1-ChR2) (Jackson Laboratories) [22] or wild type (WT) C57BL/6J mice (Charles River Laboratories) ranging from 6-10 weeks of age were used for the experiments and were randomly divided into the different experimental groups. Mice were anaesthetized with isoflurane (5% induction, 2% maintenance) during surgeries. For DRG neuronal cultures adult Thy1-ChR2 mice were used. Embryonic day 16.5 (E16.5) OF1 pregnant females were purchased from Charles River Laboratories, and the embryos were used for neuronal cortical cultures. All procedures were approved by the Ethics Committee on Animal Experimentation (CEEA) of the University of Barcelona (CEEA approval #276/16 and 141/15).

### Dorsal root ganglia (DRG) neuronal culture

DRGs from adult Thy1-ChR2 mice were dissected, washed in cooled Hank’s balanced salt solution (HBSS; ThermoFisher Scientific), and enzymatically digested (5mg/ml Dispase II (Merck) and 2.5mg/ml Collagenase Type II (ThermoFisher Scientific) in DMEM (ThermoFisher Scientific)) for 45min at 37°C. Next, the DRGs were resuspended in DMEM:F12 (ThermoFisher Scientific) media supplemented with 10% heat-inactivated Fetal Bovine Serum (FBS; ThermoFisher Scientific) and 1x B27 (ThermoFisher Scientific) and were mechanically dissociated by pipetting. After centrifugation, the resulting single cells were resuspended in culture media (DMEM:F12 media with 1x B27 and penicillin/streptomycin (P/S; ThermoFisher Scientific)) and plated in glass-coverslips (3000-4000 cells/well) or in microfluidic devices (20.000cells/device) pre-coated with 0.1mg/ml poly-D-lysine (2h, 37°C; Merck) and 2μg/ml laminin (over-night (O/N); ThermoFisher Scientific) at RT (room temperature)). An additional incubation with 5, 10 or 20 μg/ml CSPGs (Merck) was performed (2h at 37°C) in growth-inhibitory substrate experiments. Cells were allowed to grow for 24h or for 8 days (in the case of microfluidic devices) at 37°C in a 5% CO_2_ atmosphere.

### Neuronal cortical culture

E16.5 WT mice embryos were used for neuronal cortical cultures. Brains were extracted, washed in cooled 0.1M phosphate-buffered saline (PBS; ThermoFisher Scientific) containing 6.5 mg/ml glucose (Sigma Aldrich) (PBS-glucose) and the meninges excised. Both cortical lobes were dissected, sliced in a McIlwain II tissue chopper (Capdem Instruments) and trypsinized for 15 min at 37°C. The suspension was inactivated with Normal Horse Serum (NHS; ThermoFisher Scientific), incubated with 0.025% DNAse (Roche) PBS-glucose for 10 min at 37°C and mechanically dissociated. Single cells were spun down and resuspended in Neurobasal^TM^ (ThermoFisher Scientific) medium supplemented with 2mM glutamine (ThermoFisher Scientific), P/S, 6.5mg/ml glucose, NaHCO_3_ (Merck), 1x B27 and NHS 5% and plated in poly-D-lysine (0.1mg/ml) pre-coated microfluidic devices (see below) (150.000cells/device) or in glass-bottom plates (200.000cels/plate). One day after seeding, culture media was changed, and NHS was excluded from the new media. Cells were maintained at 37°C in a 5% CO2 atmosphere and culture media was changed every two days.

### Microfluidic devices

The design used for microfluidic devices were a modification of a previously published device[23, 24]. One of the devices consists in 4 reservoirs of 7mm diameter, connected in pairs by a longitudinal compartment (cell body and axonal compartments) which are in turn interconnected by 100 microchannels (10μm x 10μm x 900μm). The other used device presented a similar design and included a perpendicular channel to the microchannels, located at the center of these. The masters were produced using standard photolithography techniques at IBEC Microfab Space. Poly(dimethylsiloxane) (PDMS; Dow) was used to prepare the devices by soft lithography, which were subsequently attached to glass bottom dishes using oxygen plasma treatment.

In the first device, cortical neurons were seeded in one of the larger compartments (cell body compartment) and allowed to extend their axons across the microchannels. In the case of the second device, which was used with DRG neurons, neurons were seeded in the central channels. Vacuum aspiration in the axonal compartment allowed complete axotomies [24].

### In vitro lentiviral production and infection

LV-EF1α-hChR2(H134R)-EYFP-WPRE (LV-ChR2) produced in our laboratory were used to introduce ChR2 expression in primary cortical neurons. For LV production, 293-FT (ATCC) cells were simultaneously co-transfected with three plasmids: pMD2.G (VSV-G envelope expressing plasmid), psPAX2 (lentiviral packaging plasmid), and pLenti-EF1α-hChR2(H134R)-EYFP-WPRE (transfer plasmid) in Opti-MEM (ThermoFisher Scientific) using Lipofectamine 2000 Transfection Reagent (ThermoFisher Scientific). Transfection media was changed for culture media (Advanced DMEM (ThermoFisher Scientific) supplemented with 10% FBS, 1% P/S and 0.5% glutamine) 6h after. Media was recovered at 48h and 72h post-transfection, filtered and centrifuged at 1200g to remove debris. The supernatant was then centrifuged at 26.000rpm for 2h at 4°C in a Beckman conical tube containing a sucrose cushion (20%) for purification, ant the viral pellet was finally resuspended in PBS-1% BSA. Cortical neurons were infected at 4 days *in vitro* (DIV) for 24h, and high levels of YFP fluorescence could be observed at 7DIV, indicating positive infection.

### In vitro optogenetic stimulation

Thy1-ChR2 DRG neurons or cortical neurons infected with LV-ChR2 were used for *in vitro* optogenetic stimulation experiments. A 470nm emission LED array (LuxeonRebelTM) under the control of a Driver LED (FemtoBuck, SparkFun) of 600mA and a pulse generator PulsePal (OpenEphys)[25] was used to deliver blue light to neuronal cultures. The optogenetic stimulation protocol consisted in 1h of illumination at 20Hz of frequency with 5ms-45ms pulses, in 1s ON-1s OFF periods. In the case of DRG neurons, stimulation was applied 2h after seeding. For cortical neurons, two different stimulation time-points after the axotomy were assessed: 30 min after axotomy and 6h after axotomy. To assess neuronal activation, neurons were fixed at the end of the stimulation. Uninjured cortical neurons were stimulated in this experiment.

### In vitro chemogenetic stimulation

10µM CNO (Tocris) was added to DRG cultures 6h after axotomy and left in the media until fixation (24h after the axotomy).

### Corticospinal tract and dorsal columns axotomy

A fine incision to the skin between the shoulder blades of anaesthetized mice and subsequent muscle removal allowed thoracic vertebral column exposure. A T9 laminectomy was performed, and the dura mater was excised. In corticospinal tract injuries, two thirds of the spinal cord approximately were severed laterally with a scalpel. In dorsal column axotomies (DCA), a dorsal hemisection was conducted with fine forceps.

### Sciatic nerve crush (SNC)

The sciatic nerve was exposed after a small incision on the skin over the posterior hindlimb and blunt dissection of the gluteus maximus and the biceps femoralis. Fine forceps were used to carefully compress the nerve orthogonally 2x10s. Animals were allowed to recover for 24h, when were then sacrificed and the sciatic nerve and the sciatic DRGs (L4, L5, L6) were dissected and processed.

### In vivo optogenetic stimulation

An optic fiber cannula (1.25mm, 0.22 NA; Thorlabs) was implanted in the motor cortex (M1) of Thy1-ChR2 transgenic mice by stereotaxic surgery (–1mm antero-posterior (AP), 1.5mm lateral to Bregma, 0.5mm deep, (hind limb innervation region)) prior to injury and was fixed to the cranium with screws and dental cement. Correct placement of the cannula was tested for each animal by assessing the induction of unidirectional rotatory locomotion caused by unilateral optical stimulation of the motor component of the hind limb; animals that did not show this response were excluded from the study. At 7 days post-injury (DPI) animals started receiving daily optogenetic stimulations consisting in 1h of illumination with 470nm blue light, at 10Hz of frequency with 10ms pulses, in 1sON-4sOFF periods, for 5 consecutive days, 2 resting days followed by 5 more days. The illumination was delivered through the optic fiber cannula which was coupled to a 470nm wavelength LED source (M470F3, Thorlabs) controlled by a pulse generator (Pulse Pal) [25]. The control group went through the same procedures than the experimental group without receiving illumination.

### In vivo chemogenetic stimulation

The commercial AAV5-hSynhM3D(Gq)-mCherry or the control virus AAV5-hSyn-mCherry (Addgene) were injected into the sciatic nerve of C57BL/6J mice (1µL/sciatic nerve) 4-5 weeks before the experiment. For chemogenetic stimulation, the animals received two daily intraperitoneal (i.p.) injections of 5mg/kg CNO (Tocris). After DCA, injections were delivered starting from 3DPI and lasted 7 consecutive days. In the case of SNC, the injections were given prior to injury: from 3 days pre-injury to the same day of the injury (in total 4 days of chemogenetic stimulation).

### Behavioural assessment of sensorimotor function

Sensorimotor deficits and recovery were evaluated using the gridwalk and the BMS (Basso Mouse Scale) tests at -1, 1, 7, 14, 21, 28 and 35 DPI for CST injuries and at -1, 1, 3, 7, 10, 21 and 28 DPI for DCA. During the gridwalk animals were recorded while walking three times on a grid of 1 x 1 cm squares (total longitude of the grid: 50cm). The number of missteps in relation to the total number of steps was blindly quantified. In BMS evaluations mice were allowed to freely move in an open field and a score was assigned to each of them according to the BMS scale[26]. In brief, this scale grades from 0 to 9 the locomotor capacity of the hind limbs depending on several aspects such as ankle movement, paw placing and position or stepping.

### Tracing of injured spinal cords

A fluorescent tracer (10% Dextran-Alexa 594, 10.000MW, ThermoFisher Scientific) was injected with a stereotaxic frame (KOPF) into the motor cortex of animals with injured CST at 35DPI using the fibre optic cannula hole, in order to trace the stimulated CST. The tracer was injected at 0.2 μl/minute for a total of 1 μl, adding 5 more minutes at the end to avoid liquid spillage. 5 days were waited before sacrifice, to allow the tracer to be reach the axonal terminations

### Immunocytochemistry (ICC)

For immunocytochemistry (ICC), cells were fixed with 4% paraformaldehyde (PFA) 15 min on ice, washed and incubated in blocking solution (1% Bovine serum albumin (BSA), 0,25% Triton X-100 (0,25% Tx) in 0.1M PBS) for 1h at RT. Primary antibody (βIII tubulin (Tuj1, 1:1000, BioLegend), c-Fos (1:200, Cell Signaling), ChR2 (1:500, Progen), mCherry (1:500, Abcam) GFP (1:500, ThermoFisher Scientific) was added to the cells in blocking solution and incubated O/N at 4°C. After washing, the cells were incubated 1h at RT with Alexa-Fluor-conjugated secondary antibodies (568 goat and 488 goat and donkey) and Hoechst (Sigma Aldrich).

### Tissue processing for immunohistochemistry

Anaesthetized mice were transcardially perfused with ice-cold 4% paraformaldehyde (PFA) and the spinal cord, the brain or the DRGs were dissected and allowed to post-fix in 4% PFA for 24h at 4°C. In peripheral experiments the sciatic nerves and the DRGs were directly dissected, as perfusion was not needed, and allowed to fix in 4% PFA for 2h on ice. Fixed tissues were cryoprotected in 30% sucrose in PBS for 24h at 4°C, brains were then directly frozen and sliced at 30 μm with a freezing microtome (2000R Leica), while spinal cords and DRGs were embedded in tissue freezing medium (OCT, Sigma Aldrich), frozen and 18 μm or 10 µm slices respectively were obtained using a cryostat (CM 1900 Leica) and directly mounted. For whole mount stainings cryprotection was not needed, instead tissues were kept in 0.1M PBS.

### Immunohistochemistry (IHC)

Prior to IHC, brain slices were maintained in cryoprotection solution (30% glycerol, 30% ethylene glycol, 40% PBS). Brain IHCs were performed directly on free floating slices. Slices were blocked for 1h at RT (10% FBS, 0.5% Tx, 0.2% gelatine, 0.2M glycine in 0.1M PBS) and incubated with primary antibody O/N at 4°C (5% FBS, 0.5% Tx, 0.2% gelatine in 0.1M PBS; GFP (1:500, ThermoFisher Scientific). Slices were then repeatedly washed with PBS-0.5% Tx, incubated with secondary antibodies (Alexa Fluor 488 donkey; ThermoFisher Scientific) and Hoechst (Sigma Aldrich).

IHCs were performed directly on spinal cord or DRG preparations. Slides were blocked for 1h at RT (Blocking solution: 8% BSA, 0.3% Tx in TBS) was added and incubated for 2h at RT, followed by O/N incubation of primary antibody (GFAP (1:500, Dako); cFos (1:200, Cell Signaling); NFH (1:200; Abcam)) in 2% BSA, 0.2% Tx in TBS at 4°C. Secondary antibodies (Alexa Fluor 488 donkey and goat and 568 goat; ThermoFisher Scientific) and Hoechst were added after washing with TTBS and incubated 1h at RT.

Whole-mount IHCs were performed for sciatic nerves. Blocking solution (8% BSA, 1% Tx and 1/150 mαIgG (Fabs) in TBS) was added and incubated for 2h at RT, followed by 3 O/N of primary antibody incubation (SCG-10 (1:1000, Novus Biologicals) in 2% BSA, 0.3% Tx in TBS at 4°C. Secondary antibodies (Alexa Fluor 488 donkey) were added after washing with TTBS and incubated 1 O/N at 4°C.

Each preparation was subsequently mounted in Mowiol^TM^ (Sigma Aldrich).

### Fluorescence intensity analysis

To measure c-Fos intensity, DRG neurons were immunostained for c-Fos and ChR2 and imaged at 40x magnification with an Olympus microscope IX71 using an Orca Flash 4 (Hamamatsu Photonics). Only ChR2^+^ cells were selected for this analysis. About 20 cells per well were analysed. The nuclei of the cells were selected, and its mean c-Fos intensity determined by subtracting the corresponding background value to the integrated density (Corrected total cell fluorescence; CTCF), measured using ImageJ^TM^.

### Neurite and axonal length analysis

Images were taken at 10x magnification with an Olympus microscope IX71 using an Orca Flash 4 or a CX50 Olympus camera. TUJ1, mCherry or GFP was used to immunolabel neurites and axons. For DRG neurite analysis, three fields per well were analysed and the average neurite length per cell was obtained. Small diameter (<35µm) neurons were excluded from the analysis. In microfluidic devices, the percentage of regenerating axons normalized to the total number of axons reaching the axonal compartment was determined. For cortical neuron cultures, 8-9 fields per device were analyzed. Either average axon length or total area covered by axons was quantified. Neurite-J plugin in ImageJ^TM^ software was used for neurite and axonal measurements [27]. Area measurements were measured using ImageJ^TM^ software.

### Sprouting quantification

To quantify the number of sprouting axons before and after the lesion, we measured the tracer fluorescence intensity on sections at 0,5mm pre-injury and post-injury level. The spinal cord section images were divided in two different ROIs corresponding to grey matter and white matter for analysis.

Spinal cord sections of stimulated and non-stimulated Thy1-ChR2 mice at 500µm rostral to the lesion core were obtained using a confocal microscope (LSM 800, AxioCam 503c; Zeiss) at 10x magnification. The number of double positive axons for Dextran-Alexa 594/ChR2-YFP in the grey matter at different distances from the CST in the same section was computed and normalized to the number of Dextran-traced CST.

### Nerve regeneration analysis

Regenerating sensory axons were immunolabeled with SCG10. Whole mount preparations were imaged with a confocal microscope (LSM 800, AxioCam 503c; Zeiss) at 5x magnification. 6-8 tiles and 10-15 slices were obtained per nerve and the Blue Zen^TM^ and ImageJ^TM^ softwares were used to reconstruct the nerve and obtain a Maximum Intensity Z-projection. The crush site was determined by phase contrast images, the number of regenerating axons at several distances from the crush was determined.

### Statistical analysis

Prism 6.0 (GraphPad^TM^ Software) was used for statistics and graphical representation. Plotted data shows mean±s.e.m (standard error of the mean). Normality was determined with Shapiro-Wilk test. Significant differences are indicated by arterisks (\**p* < 0.05; \*\**p* < 0.01; \*\*\**p* < 0.005; \*\*\*\**p* < 0.001). ANOVA followed by Bonferroni post hoc test or Student’s t-Test were used in normal distributions as opposed to Mann-Whitney or Kruskal-Wallis tests as non-parametric tests for samples that did not meet normality.

## Results

### Optogenetic stimulation of DRG neurons increase their axonal growth in vitro

We first sought to determine if optogenetic activation of adult DRG neurons could enhance axonal growth *in vitro*. Adult dissociated Thy1-ChR2 DRG neurons were optically stimulated and allowed to grow for 24h.

Optical stimulation effectively increased neuronal activity in DRG neurons, as displayed by the increased levels of c-Fos staining after stimulation (49627±4373 a.u. of nuclear CTCF intensity for stimulated DRGs versus 33897±3212 a.u. of nuclear CTCF intensity for non-stimulated; *p*=0.0058 Student’s t-Test) (Fig 1A-B)

**Figure 1.**
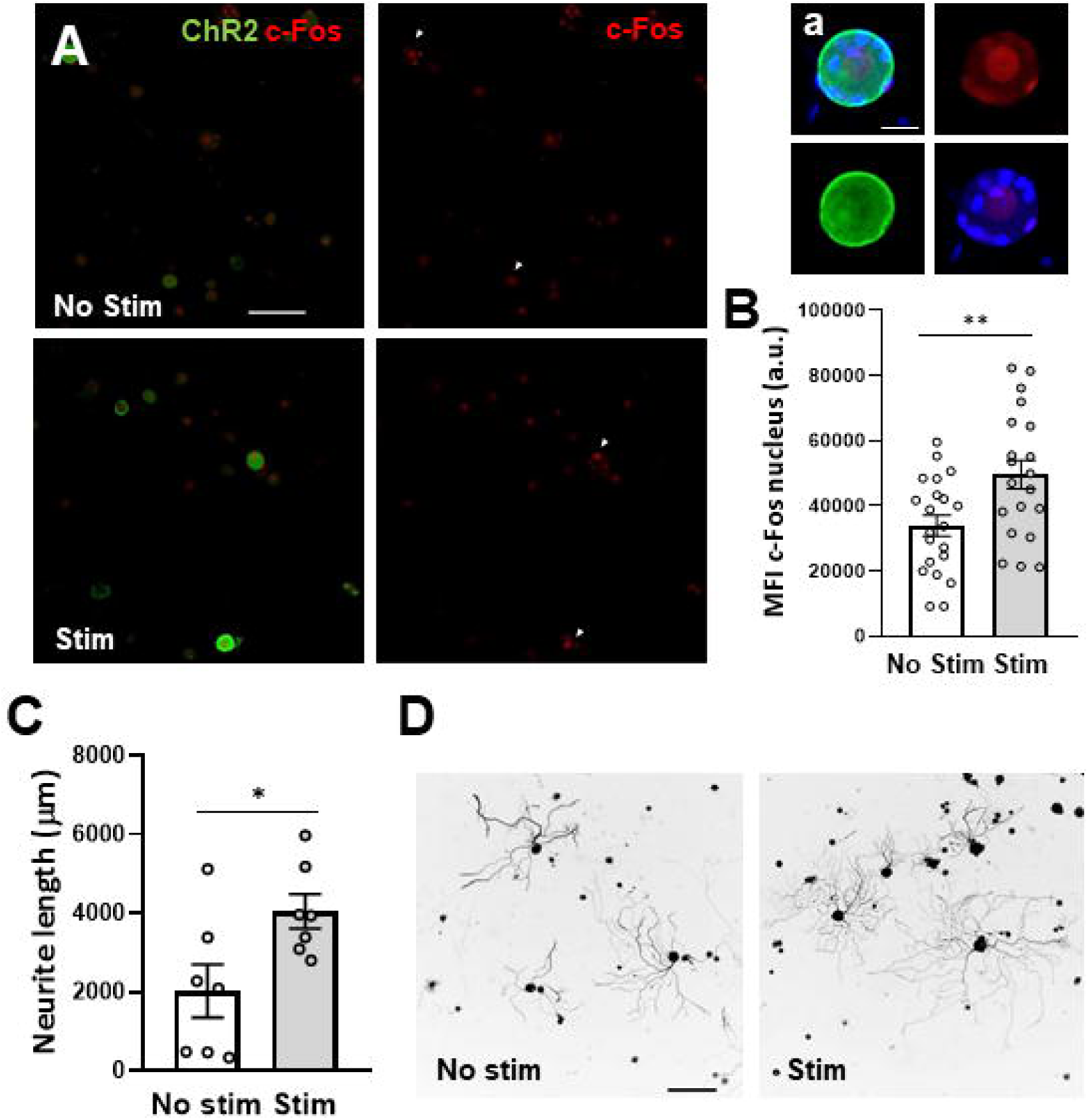
Increased neuronal activity promotes neurite outgrowth. A. Representative images of c-Fos (red) and ChR2 (green) immunostaining. White arrows depict ChR2^+^ neurons. Scale bar: 100μm. a. Higher magnification image of a stimulated neuron. ChR2 expression is mainly located in the membrane of large-diameter neurons. Scale bar: 20μm. B. c-Fos expression in the nucleus is increased just after optogenetic stimulation of Thy1-ChR2 DRG neurons (n=20-25 cells). MFI: mean fluorescence intensity. a.u.: arbitrary units. Data are expressed as mean nuclear fluorescence intensity±s.e.m. **p < 0.01 denotes significant differences in Student’s t-Test. C. Stimulated neurons presented significantly higher neurite lengths when compared to non-stimulated ones. Average neurite length per neuron was determined at 24h *in vitro*. Data are expressed as mean±s.e.m. *p < 0.05 denotes significant differences in Student’s t-Test. D. Representative images of Tuj-1 staining used to analyze neurite length. Scale bar: 200μm.

Optically stimulated neurons showed increased neurite outgrowth (4053±433.1µm) after 24h of culture when compared to non-stimulated ones (2029±647.7µm; *p*=0.0267 Student’s t-Test) (Fig 1C-D)

### Chemogenetic stimulation of DRG neurons improve their regenerative capacity after in vitro axotomy and sciatic nerve crush

We then wanted to test if this increased growth capacity also translated in regenerative capacities after axotomy. To this aim we first evaluated regenerative capacity of DRG neurons *in vitro* using microfluidic assisted axotomy and chemogenetic stimulation. DRG neurons were seeded in custom microfluidic devices as previously described, and hM3Dq or mCherry expression was induced by infection with AAVs. Axons were allowed to grow for few days through the microchannels until they reached the axonal chamber, and we then performed a vacuum assisted axotomy. CNO was administered 6 hours after axotomy. Chemogenetic stimulation resulted in axonal regeneration (*p*=0.0165; Student’s t-Test) when compared to non-stimulated mCherry controls (Fig 2A-B).

**Figure 2.**
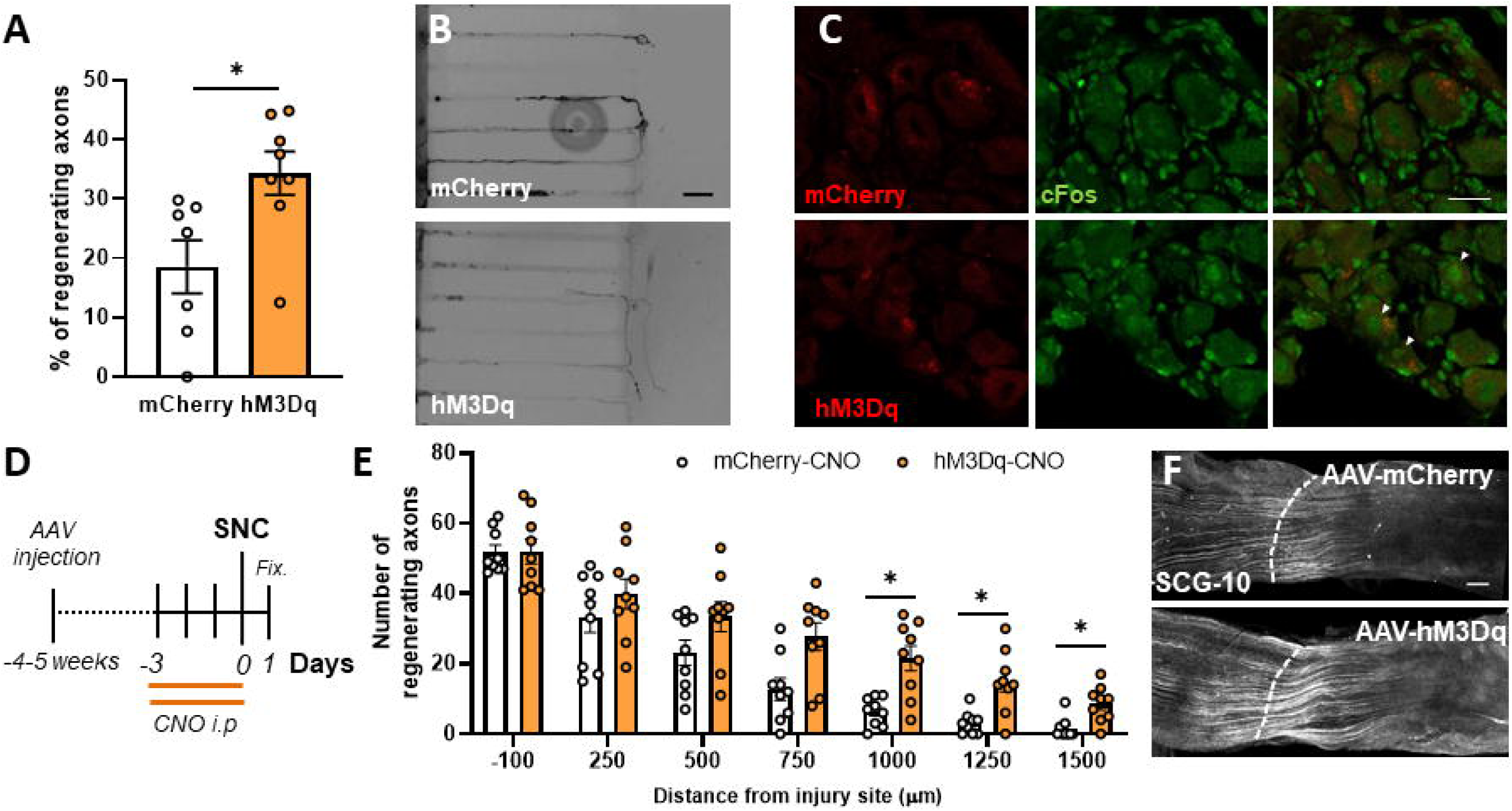
Chemogenetic stimulation induced PNS regeneration. A. Chemogenetically stimulating DRG neurons *in vitro* 6h after axotomy resulted in regeneration promotion. Data are expressed as the mean percentage of infected regenerating axons compared to the total of infected axons reaching the axonal compartment±s.e.m (n=7-8 axonal compartments). *p < 0.05 denotes significant differences in Student’s t-Test. B. Representative images of hM3Dq/mCherry^+^ axons. Scale bar: 50μm. C. c-Fos nuclear staining is observed in hM3Dq infected DRG neurons (white arrows) 1h after CNO injection, but not in mCherry^+^ DRG neurons after the same treatment. Scale bar: 25µm. D. Schematic timeline of the experiment. E. The number of regenerating sensory axons (SCG-10^+^) in stimulated nerves (hM3Dq-CNO) is increased in all assessed distances compared to the non-stimulated (mCherry-CNO) reaching statistical significance in long distances. Data are expressed as mean±s.e.m. at each distance from the injury site (n=9 sciatic nerves). *p < 0.05 denotes significant differences in ANOVA followed by Bonferroni test. F. SCG-10 immunostaining of mCherry or hM3Dq infected sciatic nerves 24h after SNC. Dotted lines indicate the injury site. Scale bar: 200μm.

We then wanted to determine if the *in vitro* results in fact translated to enhanced axon regeneration *in vivo* after PNS injury. Due to the difficulty to apply light for long periods of time in awake animals in the DRGs, we used chemogenetics for activity control. We first wanted to determine if DRGs were transduced *in vivo*. AAV5-hSyn-mCherry or AAV5-hSyn-hM3Dq-mCherry were carefully injected into the sciatic nerve and mCherry expression was examined 4-5 weeks later. Both vectors transduced DRG neurons with similar efficiency (∼35% of total DRG neurons; ∼65% of large diameter (>35µm) neurons (Supp Fig 1A-B). mCherry expression was mainly localized in the soma of large diameter NFH^+^ DRG neurons (Supp Fig 1C).

To verify the activation of DRG neurons *in vivo* after chemogenetic stimulation, we performed cFos staining 1h after CNO administration, showing increased cFos signal only in hM3Dq animals compared to mCherry controls (Fig 2B).

To assess the effects of neuronal activity on PNS regeneration, animals received 2 daily injections of 5mg/kg CNO 3 days before performing a bilateral SNC to guarantee four consecutive stimulation days (Fig 2D). 24h after injury hM3Dq stimulated animals showed increased number of regenerating sensory axons (SCG10^+^) at further distances (>750µm) when compared to mCherry controls. The two-way ANOVA showed a significant effect of the distance (*p<* 0.0001) and stimulation (*p*=0.0145) as well as their interaction (*p*=0.0101). At 750µm from the injury stimulated animals showed 27.67±4.19 axons versus 12.78±3.34 axons on non-stimulated animals (*p*=0.0694 Bonferroni post Hoc test), at 1000µm from the injury stimulated animals showed 21.56±3.72 axons versus 6.44±1.43 axons on non-stimulated animals (*p*=0.0163 Bonferroni post Hoc test), at 1250µm from the injury stimulated animals showed 14.89±3.12 axons versus 3.11±1.14 axons on non-stimulated animals (*p*=0.0261 Bonferroni post Hoc test), and at 1500µm from the injury stimulated animals showed 8.56±1.81 axons versus 1.56±1.06 axons on non-stimulated animals (*p*=0.0259 Bonferroni post Hoc test) (Fig 2D-E).

### Optogenetic stimulation of cortical neurons does not improve the sensorimotor performance after SCI

We then wanted to test if this increased regeneration capacity also was present in the CNS neurons. To this aim we first evaluated regenerative capacity of cortical neurons *in vitro* after optogenetic stimulation. Embryonic cortical neurons were seeded in custom microfluidic devices as previously described, ChR2 was induced by infection with LV-ChR2 (Fig 3A), and activation of neurons upon stimulation was assessed by cFos staining immediately after stimulation (Fig 3B). Axons were allowed to grow for few days through the microchannels until they reached the other chamber, when a vacuum assisted axotomy was performed. Optogenetic stimulation of cortical neurons decreased the axonal regrowth when stimulated 30 minutes after the axotomy Fig 3C). In contrast, 6 hours after axotomy, optogenetic stimulation resulted increase in axonal regrowth when compared to non-stimulated ones (*p*=0.0056; Mann-Whitney test; Fig 3D-F). To control the intrinsic effect of blue light on neuronal growth, we applied the light stimulation pattern on non-stimulated (GFP controls) embryonic cortical neurons and observed no difference in axonal growth (Fig 3E).

**Figure 3.**
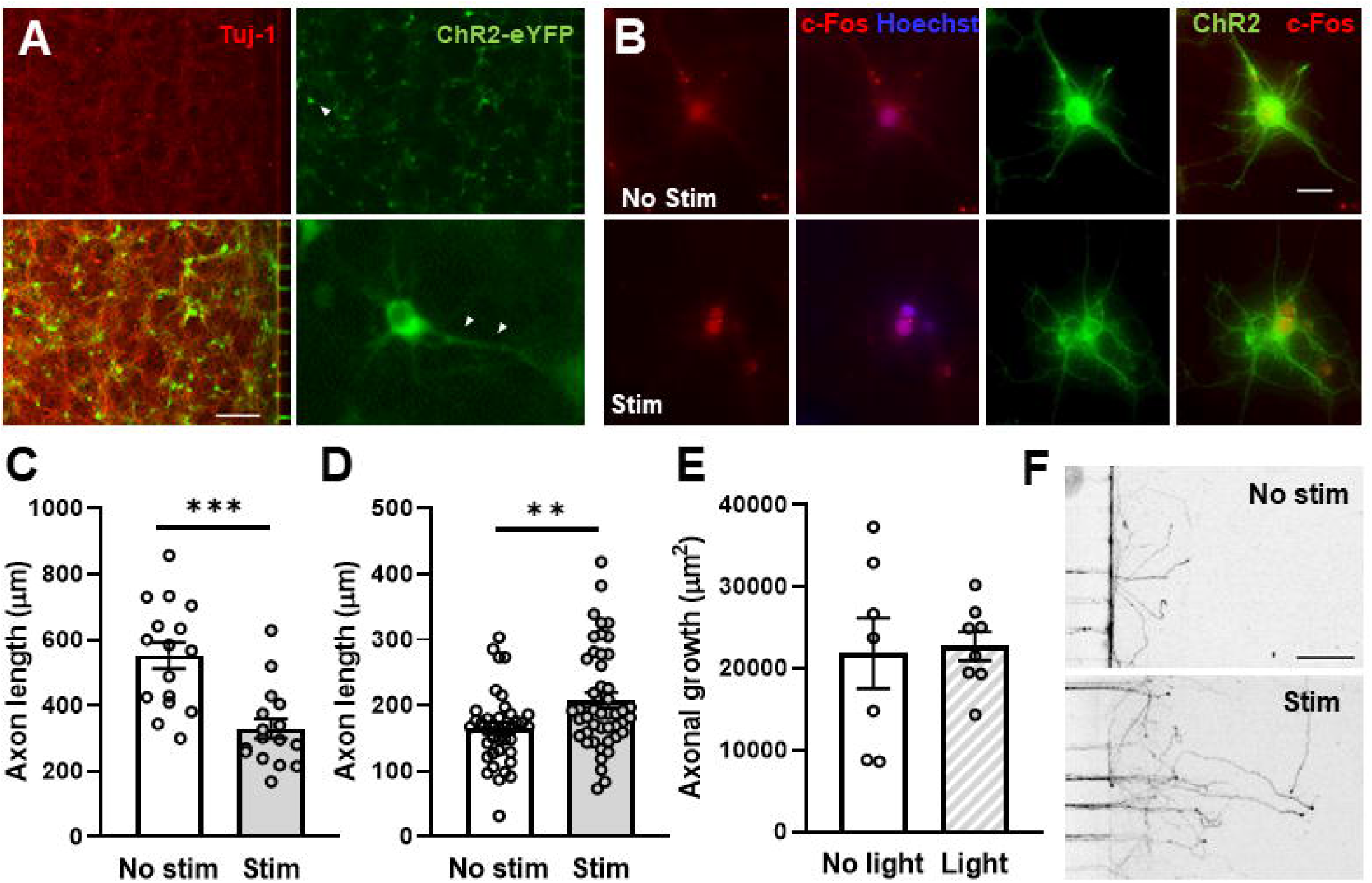
Optogenetic stimulation of cortical neurons after axotomy. A. Cortical neurons (Tuj-1) express ChR2-eYFP after LV-ChR2 infection. White arrows in the high magnification image indicate neuritic expression of ChR2. Scale bar: 250μm. B. Optogenetic stimulation increased the expression of c-Fos in cortical neurons. Representative images of c-Fos (red) and ChR2 (green) immunostaining. Scale bar: 20μm. C-D. Optogenetic stimulation of cortical neurons 30min after axotomy (B) resulted in reduced axon regeneration, while delivering the stimulation 6h after axotomy (C) increased axon regeneration. Individual ChR2^+^ axon lengths were quantified (B: n=16 images; C: n=37-43 images). Data are expressed as mean±s.e.m. ***p < 0.001 denotes significant differences in Student’s t-Test; **p < 0.01 denotes significant differences in Mann Whitney test. E. Optogenetic stimulation (Light) of WT cortical neurons did not induce any changes in axonal regeneration. Total growth area/microchannel was computed (n=7-8 images). Data are expressed as mean±s.e.m. F. Representative images of GFP/YFP staining used to analyze axon regeneration when stimulation is applied 6h after axotomy. Scale bar: 200μm.

To test if these results further translated to a better motor performance after SCI *in vivo*, we implanted optic fibre cannulas on Thy1-ChR2 animals, that have the cortical expression of ChR2 predominantly concentrated in the layer V, where corticospinal motor projecting neurons lay (Supp. Fig 2A). After recovery, animals were subjected to a CST axotomy, and optical stimulation was performed daily from day 7 after injury (Fig 4A). Stimulated animals did not show any improvement in their motor performance on the Gridwalk (Fig 4B) or the BMS (Fig 4C) when compared to non-stimulated controls. For each test evaluated, the two-way ANOVA showed a significant effect of the time (BMS *p<*0.0001; Gridwalk *p*<0.0001) but neither from the stimulation (BMS *p=*0.7201; Gridwalk *p*=0.1308) nor from their interaction (BMS *p=*0.9905; Gridwalk *p*=0.0707). In all tests, non-significant changes were observed for each timepoint when compared stimulated versus non-stimulated mice (Bonferroni multiple comparisons test).

**Figure 4.**
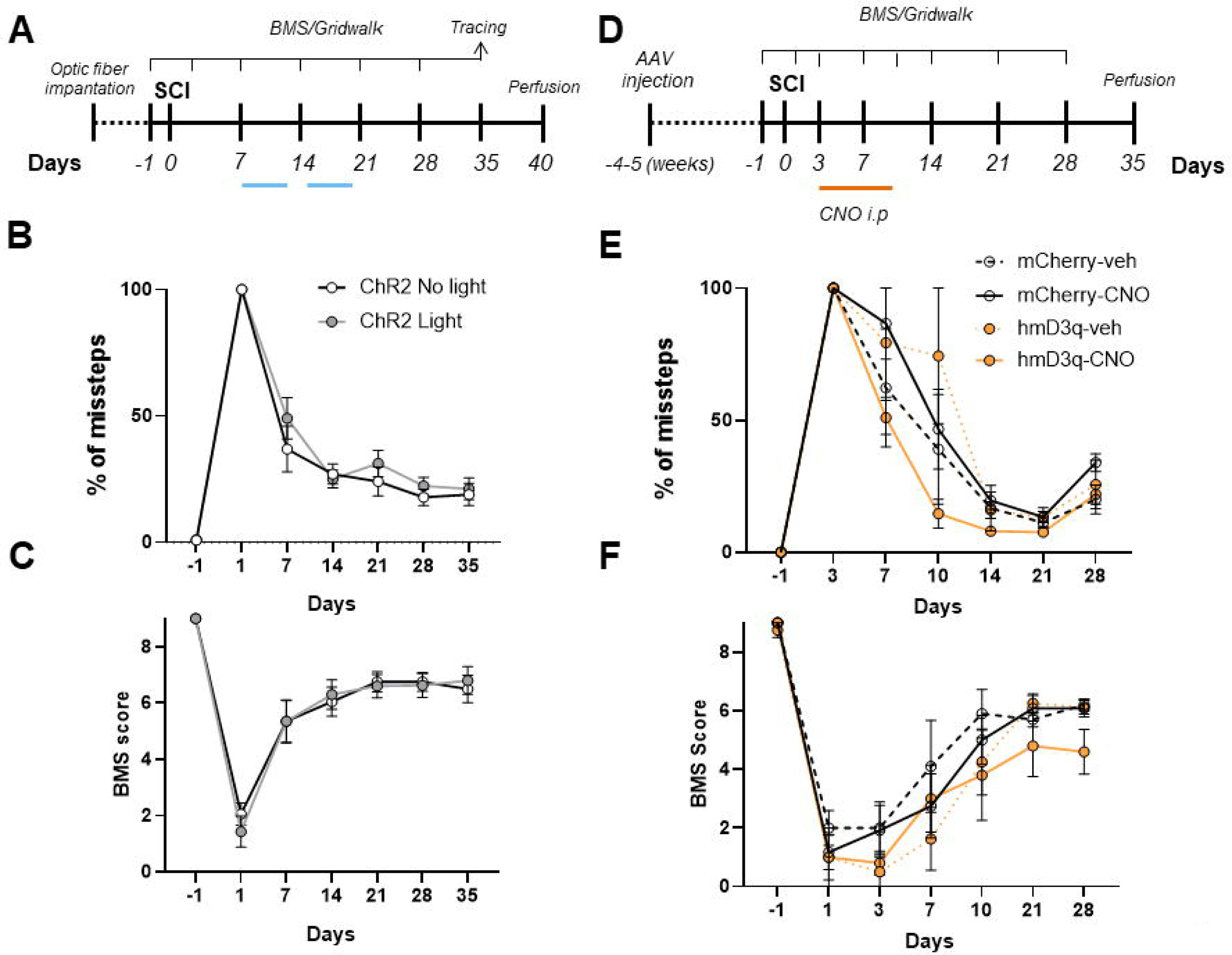
Increasing neuronal activity does not induce recovery after CNS injury. A. Timeline of the CST injury and stimulation experiments. B-C. % of missteps in the gridwalk test (B) (n=8-9 mice per group for each timepoint) and BMS score (C) (n=10 mice per group for each timepoint) show no differences in sensorimotor recovery in stimulated Thy1-ChR2 (ChR2 Light) mice when compared to non-stimulated (ChR2 No light) after CST injury. D. Timeline of the DCA and stimulation experiments. E-F. Chemogenetically stimulated animals (hM3Dq-CNO) show similar sensorimotor function recovery to non-stimulated ones (mCherry-CNO, mCherry-veh , hM3Dq-veh) after DCA as seen by the gridwalk (E) (n=3-5 mice per group for each timepoint) and BMS (F) (n=5-6 mice per group for each timepoint) tests. Represented data correspond to the BMS score and % of missteps in the gridwalk. Data are expressed as mean±s.e.m. ANOVA followed by Bonferroni post-hoctest.

These results indicate that embryonic CNS neurons have the ability to increase their axonal growth capacity *in vitro* when stimulated, but there are inhibitory signals in the CNS injury that block this regeneration *in vivo*.

### Chemogenetic stimulation of DRG neurons does not improve their sensorimotor performance after SCI

To test whether the lack of in vivo functional recovery was due to an intrinsic inability to regenerate from corticospinal neurons or the presence of an extrinsic inhibitory signal in the CNS injury, we tested the effects of chemogenetic stimulation on DRG neurons after a dorsal hemisection *in vivo.*Similarly, to what was done for the PNS injury, AAV5-hSyn-mCherry or AAV5-hSyn-hM3Dq-mCherry were carefully injected into the sciatic nerve to induce its expression on the DRG (Supp Fig 1A-C). 4-5 weeks later animals were subjected to a dorsal hemisection, and 3 days after injury animals received 2 daily injections of CNO (5mg/kg b.w.) for 7 days (Fig 4 D). As observed for the cortical stimulation, stimulated animals did not show any improvement in their motor performance on the Gridwalk (Fig 4E) or the BMS (Fig 4F) when compared to mCherry-veh, mCherry-CNO or hM3Dq-veh controls. For each test evaluated, the two-way ANOVA showed a significant effect of the time (BMS *p<*0.0001; Gridwalk *p*<0.0001) but neither from the stimulation (BMS *p=*0.2289; Gridwalk *p*=0.0694) nor from their interaction (BMS *p=*0.3614; Gridwalk *p*=0.0785). In all tests, no significant changes were observed for each timepoint when compared stimulated versus non-stimulated mice (Bonferroni multiple comparisons).

These results suggest the presence of inhibitory cues after CNS injury that could be blocking the regeneration induced by neuronal stimulation. In contrast after PNS injury where no inhibitory environment is present stimulation did induce enhanced growing capacities.***Optogenetic stimulation of DRG neurons increase their axonal growth in vitro, only on permissive substrates***

To further assess the impact of CNS inhibitory signals in the activity-induced increased axonal growth, we cultured adult DRG neurons in both permissive (Laminin) and different concentrations of inhibitory (CSPG) substrates and subjected them to optogenetic stimulation.

As previously observed (Fig 1C-D), optogenetic stimulation increased the neurite outgrowth in permissive substrates (3761±290.8µm for stimulated DRGs versus 2735±162.9µm for non-stimulated DRGs), but not on neurons seeded at any concentration of CSPGs, including the 5µg/ml concentration, that did not reduce basal growth, where optogenetic stimulation did not produce any change in neurite outgrowth (Fig 5A-B), the two-way ANOVA did not show a significant effect from the stimulation (*p*=0.4785), but it did show a significant effect of the substrate (*p*<0.0001) and their interaction ( *p*=0.0398), highlighting the blockage of the stimulation effects in the presence of CSPGs.

**Figure 5.**
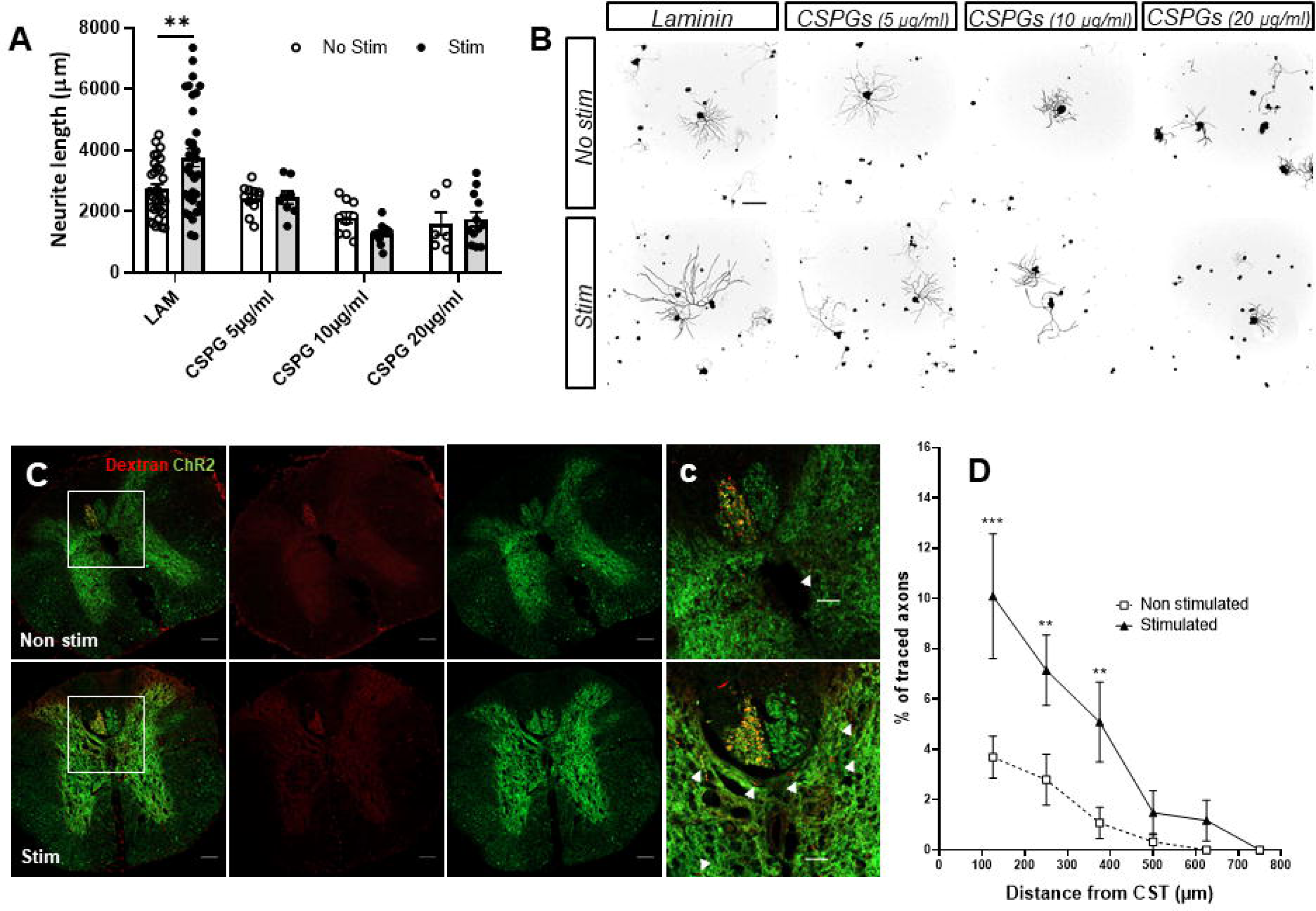
Neuronal activity induces growth but is not sufficient to overcome inhibitory environments. A. Neurite outgrowth was significantly increased in optogeneticallty stimulated DRG neurons over growth-permissive substrates (LAM: laminin), but not over different concentrations (5, 10, 20 μg/ml) of non-permissive substrates (CSPGs). Average neurite length per neuron was determined at 24h *in vitro*. Data are expressed as mean neurite length per cell±s.e.m (n=6-23 images). **p < 0.01 denotes significant differences in ANOVA followed by Bonferroni test. B. Representative images of Tuj-1 staining used to analyze neurite length. Scale bar: 200μm. C. Representative images of traced (Dextran-Alexa594) CST and ChR2-YFP in stimulated and non-stimulated Thy1-ChR2 mice in spinal cord sections 500μm rostral to the lesion core. Scale bar: 100μm. c. Higher magnification images. White arrows indicate double-positive Dextran-Alexa 594/ChR2-YFP sprouted axons. Scale bar: 50μm. D. The number of double-positive Dextran-Alexa 594/ChR2-YFP sprouted axons in the grey matter as a normalized ratio of Dextran-Alexa 594-traced CST axons shows a significantly higher percentage of sprouted axons in stimulated animals at distances up to 400µm laterally and ventrally from CST. Data are expressed as mean±s.e.m. at each distance from the CST (n=4 mice) ***p < 0.001; **p < 0.01 denote significant differences in ANOVA followed by Bonferroni test.

### Optogenetic stimulation of corticospinal motor neurons increases prelesion sprouting but does not induce regeneration across the lesion

Since activity dependant therapies have been shown to increase lateral sprouting and plasticity [17–19, 28–30], we sought to investigate if our stimulation paradigm increased the sprouting of injured axons before the lesion. To this aim, we analysed the spinal cords of stimulated Thy1-ChR2 mice after tracing their corticospinal tracts. Interestingly, we found that optogenetic stimulated animals did show more axonal tracing in the grey matter (33.60±1.13 a.u. of CTCF intensity for stimulated animals versus 23.27±0.89 a.u. of CTCF intensity on non-stimulated animals; *p*=0.0015 Student’s t-Test) in segments before (-0.5mm) the injury, but not beyond (+0.5mm) the injury (14.61±1.21 a.u. of CTCF intensity for stimulated animals versus 12.80±0.47 a.u. of CTCF intensity on non-stimulated animals; *p*=0.30328 Student’s t-Test) (Supp Fig 2B-C). Additionally, we also quantified the number of sprouted axons in the grey matter as a ratio from traced CST axons (in order to normalize the tracing efficiency between animals). Interestingly we found that stimulated animals showed a significantly higher percentage of traced sprouts up until 400µm ventrally and laterally from the CST. The two-way ANOVA showed a significant effect of the distance (*p*<0.0001), the stimulation (*p*=0.0401) as well as their interaction (*p*=0.0013). These findings suggest that the positive effects of modulation of the activity on the axonal regrowth are inhibited by CNS inhibitory signals present in the injury.

## Discussion

Neuronal activity-triggered plasticity is commonly recognized as the main component of recovery in current activity-based therapies [31, 32], however, this is mainly built on therapies that use exercise or electrical stimulation to induce neuronal activity (as in [17–19, 28–30]). Even so, even though these approaches increase neuronal activity, they do so without cellular specificity, hindering the identification of underlying cellular and molecular mechanisms induced by neuronal activity itself.

Taking advantage of the cellular specificity of optogenetics and chemogenetics [20, 21] we performed different experiments to assess the specific cellular effects of neuronal activity on axonal growth using different *in vitro* and *in vivo* neuronal models.

Consistent with previous works [23, 33–35], we showed that increasing neuronal activity on DRG sensory neurons enhanced axonal growth in non-inhibitory *in vitro* and *in vivo* peripheral injury models. Additionally, and still accordingly to previous studies [36], we also demonstrated these effects in embryonic cortical cells *in vitro,* highlighting the presence of similar mechanisms among different neuronal types.

It is important to mention that before the achievement of these results, a fine tuning of the stimulation protocol was needed, as axonal growth capacities showed to be highly dependent on the timing after injury and the pulse frequency during the stimulation (data not shown), highlighting the fact that neuronal activity needs, not only to be stimulated, but to be finely regulated in order to promote the desired outcomes. For instance, high frequency stimulations (>20Hz in 1 second trains, or >10Hz in continuous stimulation) reduced the axonal growth *in vitro,* and induced seizures *in vivo*. This has also been emphasized in previous studies indicating activity-dependant effects on gene expression [37–39] and plasticity (reviewed in [40]).

Meanwhile, timing of stimulation after injury seems to be very relevant as well, as stimulating 30 min after axotomy results in reduction of axonal growth compared to non-stimulated controls. This might be caused by the ionic imbalance generated by membrane disruption after axonal injury, that leads to exaggerated Ca^2+^ influx. Membrane sealing and restoring the ionic balance and permeability occurs early after injury, but maintaining neuronal depolarization during this period leads to the activation of some voltage gated cation channels that have been shown to inhibit growth [15, 41, 42] In accordance, little changes in the stimulation patterns might lead to opposed effects, therefore we used optogenetics whenever possible as our method of choice to stimulate activity since it allowed us a higher temporal resolution [43]. Nonetheless, the anatomical setting of the DRGs impeded the implantation of a permanent optic fiber canula, and therefore did not allow us to perform awake stimulations, as this would have compromised the wellbeing of the animals, therefore, we opted for a chemogenetic approach for *in vivo* DRG stimulation. Effectively, this method rendered similar results on neurons as those observed *in vitro* with optogenetics, resulting in enhanced regeneration. For this experiment we delivered the chemogenetic agonist (CNO) before the injury (as a preconditioning to allow enough stimulation days in this setting), leading to similar effects as those observed in previous studies using enriched environment [44] or electrical stimulation [15, 16, 45] as a preconditioning.

Since our *in vitro* and PNS injury results have shown comparable regeneration outcomes than electrical stimulations [46–48], we believe that neuronal activity itself might be responsible, at least partially, of these effects on axonal growth.

However, when we tested this paradigm in an *in vivo* model of SCI, we found that optogenetic stimulation of spinal-projecting cortical neurons did not promote the expected functional recovery after SCI, contrarily to what other studies observed using electrical stimulation [17–19], or exercise [29, 30, 49] showed.

We then focused on clarifying whether our model did not induce any kind of axonal regeneration, or it did, but was insufficient to promote recovery. In that sense, an *in vivo* CNS model of injury implies the presence of a number of factors absent in our previous experiments, including both the presence of an intrinsic lack of regenerative capacity as well as the presence of extrinsic inhibitory substrates for regeneration [9, 50, 51]. Accordingly, when we performed further histological characterization together with complementing *in vitro* experiments, we found that stimulated neurons displayed increased axonal growth in permissive substrates (laminin), but not in growth inhibitory substrates (CSPGs), and stimulated neurons showed an increased amount of growth and sprouting in the rostral vicinity of the injury, especially in the grey matter of the SC, but not caudally across or beyond the injury. While we cannot exclude that the lack of regeneration across the injury might be due, at least in part, to the intrinsic low regenerative capacity of corticospinal neurons, our data suggests that neuronal activity stimulates the axonal growth capacity of these neurons, as seen in embryonic cortical neurons *in vitro*, through a mechanism that cannot overcome the repressive signalling of growth-inhibitory molecules, such as CSPGs. Additionally, *in vivo* injured neurons when stimulated start growing through uninjured areas of the SC, similarly to what is described in other studies with classical stimulation approaches [17–19], where injured and uninjured axons grow axonal processes searching for spared intraspinal circuitry, these new connections will eventually restore the injured connections through neuronal plasticity, bypassing the injury [52]. Interestingly, classical non-specific stimulations such as epidural electrical stimulation or rehabilitation lead to stimulation of intraspinal circuits, involving spinal interneurons and motoneurons, this stimulation in turn promotes the reorganization and activation of this circuitry [53]. Whether the lack of these functional effects with our paradigm was due to a shorter stimulation period, or that it lacked the direct stimulation of this intraspinal circuitry and thus not inducing the plasticity needed, remains still unanswered. However, a recent study also observed a blockage of activity-induced axonal growth in the presence of inhibitory-substrates after cellular specific stimulation [54]. In this study, functional recovery after SCI was only achieved after combining chemogenetic stimulation and ChABC administration (a CSPG degrading enzyme) [54], surprisingly functional recovery was only achieved after 6 weeks of continuous combined treatment. In contrast to both this work and our results, both chemogenetic stimulation and visual stimulation induce functional regeneration after optic nerve crush, a CNS injury model [55]. Importantly however, the expression of the different chemogenetic receptors in this study was induced by intravitreal injection of the vectors, meaning that retinal interneurons, including amacrine and bipolar cells, will most likely be expressing the channels and be subjected to stimulation upon agonist administration. This paradigm resembles to that of electrical stimulation, considering activation of this retinal interneurons might be inducing neuronal plasticity that facilitates functional recovery.

There is plenty of evidence that neuronal activity itself is key in promoting regeneration and recovery, however, the presence of growth inhibitory substrates in the injured CNS limits the success of activity-based therapies. In line with this, combining functional rehabilitation or exercise with CSPG degrading therapies (including ChABC [56, 57] or ADAMTS4 (a disintegrin and metalloproteinase with thrombospondin motifs 4) (Griffin et al., 2020)) lead to synergistic effects, even when applied at chronic time-points [57]. These reinforce the concept that multifactorial therapies are the way to go in order to tackle the different aspects of the pathophysiology of SCIs [59].

Although neither our experiments nor other studies [54], did not demonstrate the presence of an activity-triggered transcriptional switch, more chronic stimulations could be leading to it, although with our current knowledge, more local and transient cellular mechanisms seem responsible of the the growth differences observed with stimulation of activity specifically at a cellular level. For example, neuronal activity changes the excitability of neurons through reorganization of several ionic channels [41, 60, 61]. This can in turn influence growth, as a result of intracellular ionic adjustments and their downstream associated signalling [42, 62], or even translational changes [41]. Neuronal activity can also alter neuronal metabolism, increasing glycolysis, and lipid synthesis and integration to the membrane, a cellular process essential during axonal extension [63, 64]. Another important cellular component altered by activity is the cytoskeleton and its dynamics, for instance, neuronal activity has been shown to increase dendritic spine microtubule polymerization [65]. Accordingly, chemogenetic stimulation increases microtubule dynamics in the distal axonal portion, by reducing tubulin acetylation and increasing tyrosination in this region [54]. These mechanisms may not be exclusive, on the contrary, they are likely to take place all together facilitating activity-induced axonal growth.

On the other hand, success of activity-based therapies does not depend solely on the cellular effects of activity modulations, instead, these therapies, that target activity in a more general and chronic way (as different neurons or even circuits are stimulated simultaneously), benefit from a raised general excitability that seems to be ultimate the responsible for the plasticity and reorganization that leads to recovery [52]. In that direction, rehabilitation and electrical stimulation after injury promote the formation of new synapses in the spinal cord [49, 66], respiratory function recovery after optogenetic stimulation is also attributed to synaptic plasticity [67], and a recent study showing that rehabilitation together with electrochemical stimulation promotes recovery, is credited to cortico-reticulo-spinal circuit remodelling [68]. Additionally, sprouting of spared axons, instead of injured ones, is also responsible for recovery after exercise and/or electrical stimulation [17–19, 29, 30]. In agreement to that, success of these approaches is only significant in incomplete injuries, and greater as more tissue is spared. Incomplete injuries might also help explaining why in our model we did not observe recovery after optogenetic stimulation of the motor cortex while others did [69], as different spinal injuries were used. Compellingly, compression injury leaves more uninjured tissue and spinal tracts than our injury, that axotomizes all the dorsal tracts, including the main component of the CST in mice. Besides, complete injuries also present larger glial scars, and therefore greater accumulation of growth inhibitory molecules, which translates in larger distances and hurdles to overcome or bypass to achieve functional connections, together with a greater loss of intraspinal circuits.

These observations strengthen the view that current activity-based therapies stimulate plasticity on top of inducing regeneration in specifically stimulated neurons, but probably without inducing long-lasting cellular reprogramming. This plasticity results from activity-triggered local changes, therefore prolonged stimulations are more effective increasing growth or regeneration [52]. This is also evidenced by the presence of functional recovery after 12, but not after 4 weeks of chemogenetic stimulation [54]. Therapeutically, holistic more unspecific approaches, have a more potent effect than cellular specific stimulations alone, as evidenced by enriched environment compared to chemogenetic stimulation, which presents a lower rate of regeneration [44]. Studies have also shown that this activity-induced plasticity can be accelerated and improved by linking the activation of different relays topographically in a system [53, 70, 71]. These systems are however more challenging to define underlying mechanisms and understand the true nature of the gains of these therapies.

Briefly, we found that specific cellular stimulation of neuronal activity induced axonal growth, but only in the absence of inhibitory substrates. Likewise, this approach seems to be therapeutically less efficient in enhancing recovery than other more chronic or general stimulations. This also suggests and strengthens the idea that activity-based therapies succeed because of local transient cellular changes coupled with neuronal plasticity, rather than resulting in a cellular reprogramming of growth capacities, and because of that, longer stimulation periods elicit more robust responses.

## Conclusions

**List of abbreviations**

**Declarations**

### Ethics approval and consent to participate

Not applicable

### Consent for publication

Not applicable

### Availability of data and materials

All data generated or analysed during this study are included in this published article or available from the corresponding author on reasonable request.

## Funding

This research was supported by HDAC3-EAE-SCI Project with ref. PID2020-119769RA-I00 from MCIN/ AEI /10.13039/501100011033 to AH and PRPSEM Project with ref. RTI2018-099773-B-I00 from MCINN/AEI/10.13039/501100011033/ FEDER “Una manera de hacer Europa”, the CERCA Programme, and the Commission for Universities and Research of the Department of Innovation, Universities, and Enterprise of the Generalitat de Catalunya (SGR2017-648) to JADR. The project leading to these results received funding from the María de Maeztu Unit of Excellence (Institute of Neurosciences, University of Barcelona) MDM-2017-0729 and Severo Ochoa Unit of Excellence (Institute of Bioengineering of Catalonia) CEX2018-000789-S from MCIN/AEI /10.13039/501100011033. FMV was supported by a fellowship from the “Ayudas para la Formación de Profesorado Universitario” (FPU16/03992) program, from the Spanish Ministry of Universities.

### Authors’ contributions

FMV performed, designed experiments, performed data analysis, and wrote the manuscript; SMT performed and designed experiments and performed data analysis; JADR supervised experiments, provided experimental funds and edited the manuscript; AH performed, designed experiments, performed data analysis, provided experimental funds, and wrote the manuscript.

## Acknowledgements

The authors thank Miriam Segura-Feliu and Ana López-Mengual for their technical help.

---

**Supplementary Figure 1. AAV-mCherry and AAV-hM3Dq infection of DRG neurrons.** A. Images of mCherry expression in AAV-mCherry and AAV-hM3Dq infected DRG neurons 4-5 weeks after viral injection. Scale bar: 100μm. B. Large-diameter DRG neurons are preferentially infected. % of large-diameter (diameter >35μm) infected neurons was analyzed (n=7-9 DRGs). Data are expressed as mean±s.e.m. C. Both AAVs infect NFH^+^ cells. Scale bar: 30μm.

**Supplementary Figure 2. ChR2 expression and growth after CST axotomy in Thy1-ChR2 mice.** A. While some layer II-III neurons express ChR2, expression is concentrated in layer V neurons. Scale bar: 100μm. B. Schematic representation of the quantification in C. The spinal cord was divided in grey matter and white matter for analysis. C. Tracer intensity quantification of injured stimulated and non-stimulated Thy1-ChR2 mice at 0,5mm pre-injury and post-injury. Optogenetic stimulation of the CST resulted in increased sprouting in the grey matter pre-injury, but not post-injury. a.u.: arbitrary units. Data are expressed as mean tracer intensity±s.e.m (n=5 spinal cords). **p < 0.01 denotes significant differences in Student’s t-Test.

## References

1. Fitch MT, Silver J (2008) CNS injury, glial scars, and inflammation: Inhibitory extracellular matrices and regeneration failure. Experimental Neurology. https://doi.org/10.1016/j.expneurol.2007.05.014

2. Richardson PM, McGuinness UM, Aguayo AJ (1980) Axons from CNS neurones regenerate into PNS grafts. Nature. https://doi.org/10.1038/284264a0

3. Huebner EA, Strittmatter SM (2009) Axon regeneration in the peripheral and central nervous systems. Results and Problems in Cell Differentiation. https://doi.org/10.1007/400_2009_19

4. Cregg JM, DePaul MA, Filous AR, Lang BT, Tran A, Silver J (2014) Functional regeneration beyond the glial scar. Experimental Neurology. https://doi.org/10.1016/j.expneurol.2013.12.024

5. Jones LL, Sajed D, Tuszynski MH (2003) Axonal Regeneration through Regions of Chondroitin Sulfate Proteoglycan Deposition after Spinal Cord Injury: A Balance of Permissiveness and Inhibition. Journal of Neuroscience. https://doi.org/10.1523/jneurosci.23-28-09276.2003

6. Ohtake Y, Li S (2015) Molecular mechanisms of scar-sourced axon growth inhibitors. Brain Research. https://doi.org/10.1016/j.brainres.2014.08.064

7. Sami A, Selzer ME, Li S (2020) Advances in the Signaling Pathways Downstream of Glial-Scar Axon Growth Inhibitors. Frontiers in Cellular Neuroscience. https://doi.org/10.3389/fncel.2020.00174

8. Curcio M, Bradke F (2018) Axon Regeneration in the Central Nervous System: Facing the Challenges from the Inside. Annual Review of Cell and Developmental Biology. https://doi.org/10.1146/annurev-cellbio-100617-062508

9. Mahar M, Cavalli V (2018) Intrinsic mechanisms of neuronal axon regeneration. Nature Reviews Neuroscience. https://doi.org/10.1038/s41583-018-0001-8

10. He Z, Jin Y (2016) Intrinsic Control of Axon Regeneration. Neuron. https://doi.org/10.1016/j.neuron.2016.04.022

11. Anderson MA, O’Shea TM, Burda JE, et al (2018) Required growth facilitators propel axon regeneration across complete spinal cord injury. Nature. https://doi.org/10.1038/s41586-018-0467-6

12. Wu D, Klaw MC, Connors T, Kholodilov N, Burke RE, Tom VJ (2015) Expressing constitutively active rheb in adult neurons after a complete spinal cord injury enhances axonal regeneration beyond a chondroitinase-treated glial scar. Journal of Neuroscience. https://doi.org/10.1523/JNEUROSCI.0719-15.2015

13. Arlotta P, Molyneaux BJ, Chen J, Inoue J, Kominami R, MacKlis JD (2005) Neuronal subtype-specific genes that control corticospinal motor neuron development in vivo. Neuron 45:207–221

14. Friel KM, Williams PTJA, Serradj N, Chakrabarty S, Martin JH (2014) Activity-Based Therapies for Repair of the Corticospinal System Injured during Development. Frontiers in Neurology. https://doi.org/10.3389/fneur.2014.00229

15. Goganau I, Sandner B, Weidner N, Fouad K, Blesch A (2018) Depolarization and electrical stimulation enhance in vitro and in vivo sensory axon growth after spinal cord injury. Experimental Neurology 300:247–258

16. Udina E, Furey M, Busch S, Silver J, Gordon T, Fouad K (2008) Electrical stimulation of intact peripheral sensory axons in rats promotes outgrowth of their central projections. Experimental Neurology 210:238–247

17. Carmel JB, Kimura H, Berrol LJ, Martin JH (2013) Motor cortex electrical stimulation promotes axon outgrowth to brain stem and spinal targets that control the forelimb impaired by unilateral corticospinal injury. European Journal of Neuroscience 37:1090–1102

18. Carmel JB, Kimura H, Martin JH (2014) Electrical stimulation of motor cortex in the uninjured hemisphere after chronic unilateral injury promotes recovery of skilled locomotion through ipsilateral control. Journal of Neuroscience 34:462–466

19. Carmel JB, Berrol LJ, Brus-Ramer M, Martin JH (2010) Chronic electrical stimulation of the intact corticospinal system after unilateral injury restores skilled locomotor control and promotes spinal axon outgrowth. Journal of Neuroscience 30:10918–10926

20. Deisseroth K (2015) H I S TO R I C A L C O M M E N TA RY Optogenetics : 10 years of microbial opsins in neuroscience. 18:1213–1225

21. Sternson SM, Roth BL (2014) Chemogenetic tools to interrogate brain functions. Annual Review of Neuroscience 37:387–407

22. Arenkiel BR, Peca J, Davison IG, Feliciano C, Deisseroth K, Augustine GJJ, Ehlers MD, Feng G (2007) In Vivo Light-Induced Activation of Neural Circuitry in Transgenic Mice Expressing Channelrhodopsin-2. Neuron. https://doi.org/10.1016/j.neuron.2007.03.005

23. Sala-Jarque J, Mesquida-Veny F, Badiola-Mateos M, Samitier J, Hervera A, del Río JA (2020) Neuromuscular Activity Induces Paracrine Signaling and Triggers Axonal Regrowth after Injury in Microfluidic Lab-On-Chip Devices. Cells 9:302

24. Taylor AM, Blurton-Jones M, Rhee SW, Cribbs DH, Cotman CW, Jeon NL (2005) A microfluidic culture platform for CNS axonal injury, regeneration and transport. Nature Methods. https://doi.org/10.1038/nmeth777

25. Siegle JH, López AC, Patel YA, Abramov K, Ohayon S, Voigts J (2017) Open Ephys: An open-source, plugin-based platform for multichannel electrophysiology. Journal of Neural Engineering. https://doi.org/10.1088/1741-2552/aa5eea

26. Basso DM, Fisher LC, Anderson AJ, Jakeman LB, McTigue DM, Popovich PG (2006) Basso mouse scale for locomotion detects differences in recovery after spinal cord injury in five common mouse strains. Journal of Neurotrauma. https://doi.org/10.1089/neu.2006.23.635

27. Meijering E, Jacob M, Sarria JCF, Steiner P, Hirling H, Unser M (2004) Design and Validation of a Tool for Neurite Tracing and Analysis in Fluorescence Microscopy Images. Cytometry Part A. https://doi.org/10.1002/cyto.a.20022

28. Sánchez-Ventura J, Giménez-Llort L, Penas C, Udina E (2021) Voluntary wheel running preserves lumbar perineuronal nets, enhances motor functions and prevents hyperreflexia after spinal cord injury. Experimental Neurology. https://doi.org/10.1016/j.expneurol.2020.113533

29. Goldshmit Y, Lythgo N, Galea MP, Turnley AM (2008) Treadmill training after spinal cord hemisection in mice promotes axonal sprouting and synapse formation and improves motor recovery. Journal of Neurotrauma 25:449–465

30. Engesser-Cesar C, Ichiyama RM, Nefas AL, Hill MA, Edgerton VR, Cotman CW, Anderson AJ (2007) Wheel running following spinal cord injury improves locomotor recovery and stimulates serotonergic fiber growth. Eur J Neurosci 25:1931–1939

31. Carulli D, Foscarin S, Rossi F (2011) Activity-Dependent Plasticity and Gene Expression Modifications in the Adult CNS. Frontiers in Molecular Neuroscience 4:1–12

32. Hogan MK, Hamilton GF, Horner PJ (2020) Neural Stimulation and Molecular Mechanisms of Plasticity and Regeneration: A Review. Frontiers in Cellular Neuroscience 14:1–16

33. Jaiswal PB, English AW (2017) Chemogenetic enhancement of functional recovery after a sciatic nerve injury. European Journal of Neuroscience 45:1252–1257

34. Ward PJ, Jones LN, Mulligan A, Goolsby W, Wilhelm JC, English AW (2016) Optically-induced neuronal activity is sufficient to promote functional motor axon regeneration in vivo. PLoS ONE 11:1–16

35. Park S, Koppes RA, Froriep UP, Jia X, Achyuta AKH, McLaughlin BL, Anikeeva P (2015) Optogenetic control of nerve growth. Scientific Reports 5:1–9

36. Ward PJ, Clanton SL, English AW (2018) Optogenetically enhanced axon regeneration: motor versus sensory neuron-specific stimulation. European Journal of Neuroscience. https://doi.org/10.1111/ejn.13836

37. Tyssowski KM, DeStefino NR, Cho JH, et al (2018) Different Neuronal Activity Patterns Induce Different Gene Expression Programs. Neuron 98:530–546.e11

38. Lee PR, Cohen JE, Iacobas DA, Iacobas S, Douglas Fields R (2017) Gene networks activated by specific patterns of action potentials in dorsal root ganglia neurons. Scientific Reports 7:1– 14

39. Miyasaka Y, Yamamoto N (2021) Neuronal Activity Patterns Regulate Brain-Derived Neurotrophic Factor Expression in Cortical Cells via Neuronal Circuits. Frontiers in Neuroscience. https://doi.org/10.3389/fnins.2021.699583

40. Fauth M, Tetzlaff C (2016) Opposing effects of neuronal activity on structural plasticity. Frontiers in Neuroanatomy 10:1–18

41. Enes J, Langwieser N, Ruschel J, et al (2010) Electrical activity suppresses axon growth through Cav1.2 channels in adult primary sensory neurons. Current Biology 20:1154–1164

42. Tedeschi A, Dupraz S, Laskowski CJ, Xue J, Ulas T, Beyer M, Schultze JL, Bradke F (2016) The Calcium Channel Subunit Alpha2delta2 Suppresses Axon Regeneration in the Adult CNS. Neuron 92:419–434

43. Rost BR, Schneider-warme F, Schmitz D, Hegemann P (2017) Primer Optogenetic Tools for Subcellular Applications in Neuroscience. Neuron 96:572–603

44. Hutson TH, Kathe C, Palmisano I, et al (2019) Cbp-dependent histone acetylation mediates axon regeneration induced by environmental enrichment in rodent spinal cord injury models. Science Translational Medicine. https://doi.org/10.1126/scitranslmed.aaw2064

45. Senger JLB, Verge VMK, Macandili HSJ, Olson JL, Chan KM, Webber CA (2018) Electrical stimulation as a conditioning strategy for promoting and accelerating peripheral nerve regeneration. Experimental Neurology 302:75–84

46. Ahlborn P, Schachner M, Irintchev A (2007) One hour electrical stimulation accelerates functional recovery after femoral nerve repair. Experimental Neurology 208:137–144

47. Al-Majed AA, Siu LT, Gordon T (2004) Electrical stimulation accelerates and enhances expression of regeneration-associated genes in regenerating rat femoral motoneurons. Cellular and Molecular Neurobiology 24:379–402

48. Al-Majed AA, Brushart TM, Gordon T (2000) Electrical stimulation accelerates and increases expression of BDNF and trkB mRNA in regenerating rat femoral motoneurons. Eur J Neurosci 12:4381–4390

49. Loy K, Schmalz A, Hoche T, Jacobi A, Kreutzfeldt M, Merkler D, Bareyre FM (2018) Enhanced Voluntary Exercise Improves Functional Recovery following Spinal Cord Injury by Impacting the Local Neuroglial Injury Response and Supporting the Rewiring of Supraspinal Circuits. J Neurotrauma 35:294–2915

50. Bradbury EJ, Burnside ER (2019) Moving beyond the glial scar for spinal cord repair. Nature Communications. https://doi.org/10.1038/s41467-019-11707-7

51. Mesquida-Veny F, del Río JA, Hervera A (2021) Macrophagic and microglial complexity after neuronal injury. Progress in Neurobiology. https://doi.org/10.1016/j.pneurobio.2020.101970

52. Courtine G, Sofroniew M v. (2019) Spinal cord repair: advances in biology and technology. Nature Medicine. https://doi.org/10.1038/s41591-019-0475-6

53. Wagner FB, Mignardot JB, le Goff-Mignardot CG, et al (2018) Targeted neurotechnology restores walking in humans with spinal cord injury. Nature 563:65–93

54. Wu D, Jin Y, Shapiro TM, Hinduja A, Baas PW, Tom VJ (2020) Chronic neuronal activation increases dynamic microtubules to enhance functional axon regeneration after dorsal root crush injury. Nature Communications 11:1–16

55. Lim JHA, Stafford BK, Nguyen PL, Lien B V., Wang C, Zukor K, He Z, Huberman AD (2016) Neural activity promotes long-distance, target-specific regeneration of adult retinal axons. Nature Neuroscience 19:1073–1084

56. García-Alías G, Barkhuysen S, Buckle M, Fawcett JW (2009) Chondroitinase ABC treatment opens a window of opportunity for task-specific rehabilitation. Nature Neuroscience 12:1145– 1151

57. Wang D, Ichiyama RM, Zhao R, Andrews MR, Fawcett JW (2011) Chondroitinase combined with rehabilitation promotes recovery of forelimb function in rats with chronic spinal cord injury. 31:9332–9344

58. Griffin JM, Fackelmeier B, Clemett CA, Fong DM, Mouravlev A, Young D, O’Carroll SJ (2020) Astrocyte-selective AAV-ADAMTS4 gene therapy combined with hindlimb rehabilitation promotes functional recovery after spinal cord injury. Experimental Neurology. https://doi.org/10.1016/j.expneurol.2020.113232

59. Griffin JM, Bradke F (2020) Therapeutic repair for spinal cord injury: combinatory approaches to address a multifaceted problem. EMBO Molecular Medicine 12:1–29

60. Misonou H, Mohapatra DP, Park EW, Leung V, Zhen D, Misonou K, Anderson AE, Trimmer JS (2004) Regulation of ion channel localization and phosphorylation by neuronal activity. Nature Neuroscience 7:711–718

61. Romer SH, Deardorff AS, Fyffe REW (2016) Activity-dependent redistribution of Kv2.1 ion channels on rat spinal motoneurons. Physiological Reports 4:1–11

62. Shim S, Ming GL (2010) Roles of channels and receptors in the growth cone during PNS axonal regeneration. Experimental Neurology 223:38–44

63. Bas-Orth C, Tan YW, Lau D, Bading H (2017) Synaptic activity drives a genomic program that promotes a neuronal warburg effect. Journal of Biological Chemistry 292:5183–5194

64. Segarra-Mondejar M, Casellas-Díazi S, Ramiro-Paretai M, Müller-Sánchez C, Martorell-Riera A, Hermelo I, Reina M, Aragonés J, Martínez-Estrada OM, Soriano FX (2018) Synaptic activity-induced glycolysis facilitates membrane lipid provision and neurite outgrowth. The EMBO Journal 37:1–16

65. Hu X, Viesselmann C, Nam S, Merriam E, Dent EW (2008) Activity-dependent dynamic microtubule invasion of dendritic spines. Journal of Neuroscience 28:13094–13105

66. Eisdorfer JT, Smit RD, Keefe KM, Lemay MA, Smith GM, Spence AJ (2020) Epidural Electrical Stimulation: A Review of Plasticity Mechanisms That Are Hypothesized to Underlie Enhanced Recovery From Spinal Cord Injury With Stimulation. Frontiers in Molecular Neuroscience 13:1–12

67. Alilain WJ, Li X, Horn KP, Dhingra R, Dick TE, Herlitze S, Silver J (2008) Light-induced rescue of breathing after spinal cord injury. Journal of Neuroscience 28:11862–11870

68. Asboth L, Friedli L, Beauparlant J, et al (2018) Cortico-reticulo-spinal circuit reorganization enables functional recovery after severe spinal cord contusion. Nat Neurosci 21:576–588

69. Deng WW, Wu GY, Min LX, Feng Z, Chen H, Tan ML, Sui JF, Liu HL, Hou JM (2021) Optogenetic Neuronal Stimulation Promotes Functional Recovery After Spinal Cord Injury. Frontiers in Neuroscience 15:1–10

70. McPherson JG, Miller RR, Perlmutter SI, Poo MM (2015) Targeted, activity-dependent spinal stimulation produces long-lasting motor recovery in chronic cervical spinal cord injury. Proc Natl Acad Sci U S A 112:12193–12198

71. Mishra AM, Pal A, Gupta D, Carmel JB (2017) Paired motor cortex and cervical epidural electrical stimulation timed to converge in the spinal cord promotes lasting increases in motor responses. Journal of Physiology 595:6953–6968

