## Supplementary figures and images for "Genetic control of neuronal activity enhances axonal growth only on permissive substrates"

### Supplementary figure 1

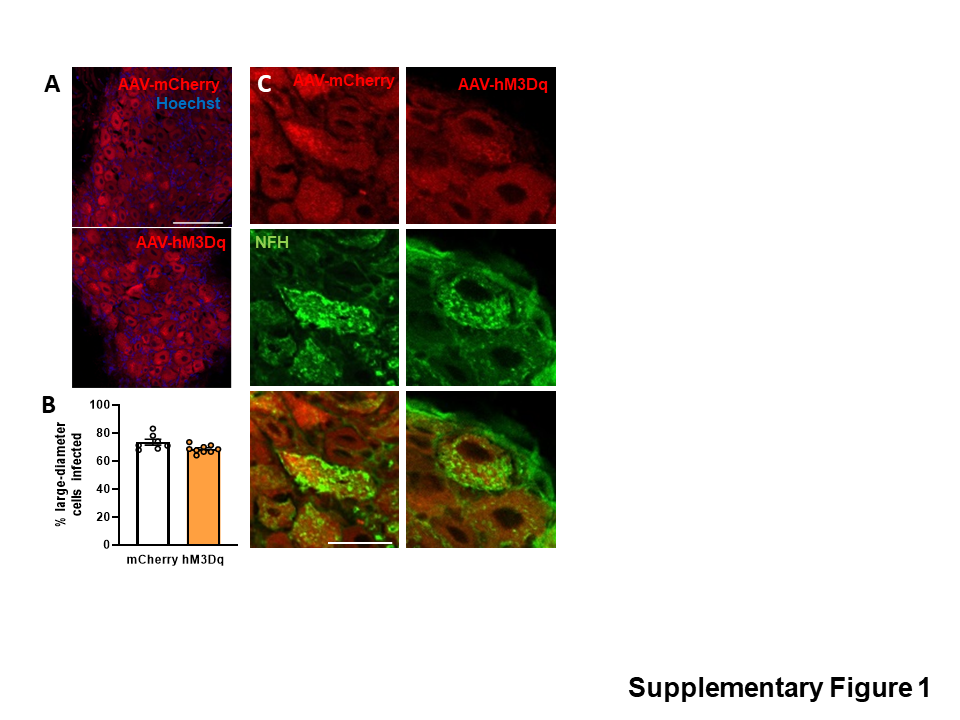

### Supplementary figure 2

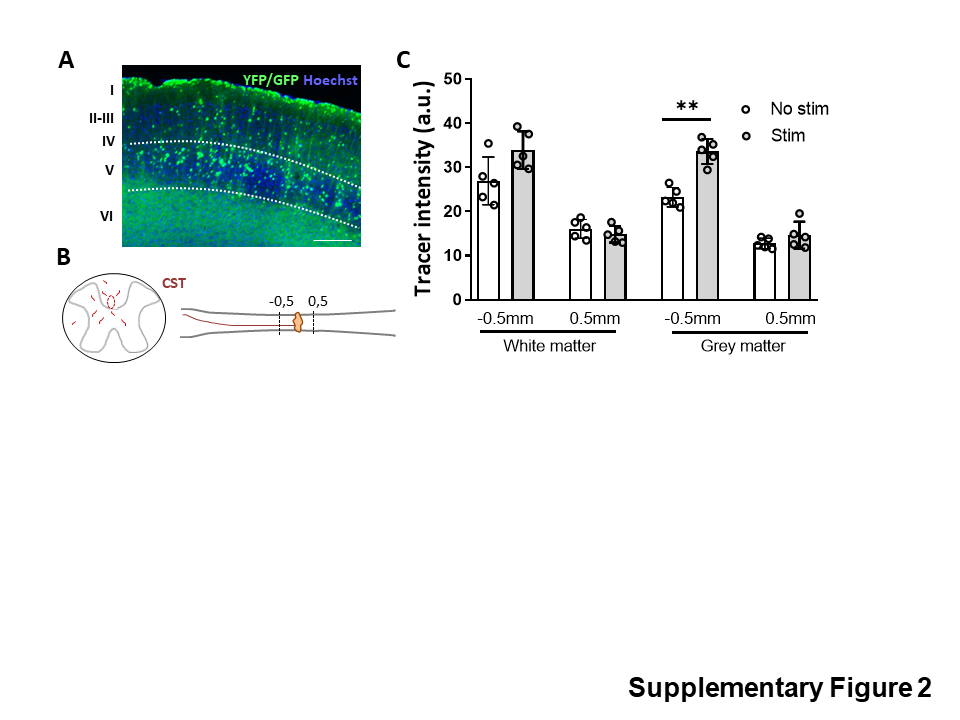
